# Accelerating Inflammation Resolution to Counteract Chemical Cutaneous Injury

**DOI:** 10.1101/749184

**Authors:** Satyanarayana Achanta, Narendranath Reddy Chintagari, Shrilatha Balakrishna, Boyi Liu, Sven-Eric Jordt

## Abstract

Chemical exposure to vesicants such as sulfur mustard (SM), and electrophilic riot control agents such as 2-chlorobenzalmalononitrile (CS) tear gas agent, cause strong cutaneous inflammation. Classical anti-inflammatory treatments have focused on interference with target initiation and maintenance of inflammation, with mixed outcomes. Inflammation is broadly classified into three temporal phases, initiation, amplification and maintenance, and resolution. Resolution of inflammation was thought to be a passive process but the recent body of literature shows that resolution is an active process and is mediated by fatty acid-derived mediators (specialized pro-resolving mediators, SPMs). We hypothesized that accelerating resolution phase of inflammation may attenuate the exaggerated inflammatory response following chemical threat exposure, leading to decreased morbidity and improved recovery. In this study, SPMs, such as Resolvin D1 (RvD1) and Resolvin D2 (RvD2), were administered to mice at nanogram doses post-exposure to an SM analog, 2-chloroethyl-ethyl-sulfide (CEES) or CS tear gas agent. SPMs decreased edema (ear thickness and punch biopsy weights), pro-inflammatory cytokines (IL-1β, CXCL1/KC, MIP2) and protease marker (MMP-9), and vascular leakage (determined by IRDye 800 CW PEG) while improving histopathology in cutaneous chemical injury mouse models. These results support our hypothesis and pave the way for SPMs for further development as potential medical countermeasures for chemical threat agents-induced skin injuries.

## Introduction

### Chemical threat agents-induced skin injuries

The treatment of injuries caused by exposures to chemical warfare agents or hazardous industrial chemicals remains challenging due to limited information about the injury mechanisms and biological targets. Exposures to chemical threat agents oftentimes induce strong inflammatory tissue responses that contribute to morbidity and delayed tissue repair and recovery. Skin blistering agents (vesicants) such as sulfur mustard (HD or SM) in mustard gas, or its analog, 2-chloroethyl-ethyl-sulfide (CEES), while not immediately irritating, induce delayed inflammatory responses that contribute to the separation of skin layers and blister formation hours after exposure (1-6). This response is initiated by the alkylating activity of HD and CEES, followed by induction of oxidative stress and the formation of lipid peroxidation products. Within a short time, pro-inflammatory cytokines and chemokines can be detected, including IL-1α/β, IL-6, IL-8, TNF-α, GM-CSF and KC, and inflammatory cells infiltrate the exposed skin, including neutrophils, mast cells, and macrophages. Proteases such as MMP-9 are induced that contribute to edema formation, tissue destruction and blistering (6). Sulfur mustard wound often shows incomplete healing, with patients burdened by chronic skin inflammation and pruritus even decades after exposure (2, 7).

Other chemical agents cause more immediate harm, followed by an exaggerated inflammatory response that aggravates tissue injury and delays recovery. For example, skin and eye exposures to tear gas agents can cause immediate chemical burns and irritation due to their high chemical reactivity leading to the rapid modification of proteins, lipids and other biomolecules (8). Skin exposure to tear gas agents such as 2-chlorobenzalmalononitrile (CS tear gas agent) cause pain and irritation, and, depending on the exposure concentration and site, can lead to burns, subcutaneous edema and bulla formation (8-11). Some studies have reported contact dermatitis following repeated exposure to CS tear gas agent (12-14). Several short- and long-term effects of exposure to CS tear gas agent were reported (14-16).

While inflammation is essential for host defense and may prevent infiltration of the chemical wound by pathogens and contribute to clearance of debris, experiments in chemically exposed neutrophil-depleted animals demonstrated that these pro-inflammatory cells exacerbate the injury and contribute to morbidity (17). Taken together, these reports support the idea that exaggerated inflammation is a major driver of progressive tissue injury hours (and even days) after the chemical threat exposure has ended.

Several treatment options, including the use of nonsteroidal anti-inflammatory drugs (NSAIDs), are published for HD and CS tear gas-induced cutaneous injury, with mixed outcomes (8, 18, 19). Despite their use as chemical warfare agents more than a century, there is no specific antidote for sulfur mustard or tear gas agent-induced cutaneous injuries (9). Although some therapeutic agents were tested for SM and CS tear agent-induced injuries, symptomatic treatment is the general line of treatment (8, 9, 18, 20-23). Decontamination of affected subjects is crucial in mitigating the injuries. However, decontamination of contaminated surfaces is also difficult as CS tear gas agent is almost insoluble in water and only slightly soluble in ethyl alcohol and carbon tetrachloride. Similarly, the presence of sulfur mustard in different physical states complicates decontamination. The actual concentrations of incapacitating agents exposed to civilians and law enforcement authorities are highly variable, which further complicates the triaging of clinical symptoms and medical management. While some of the potential therapeutic drug candidates are being tested in pre-clinical studies and are promising, none of the compounds were approved by US FDA (6, 8, 18, 19, 24-28). The lack of effective antidotes for SM and CS tear gas-induced skin injuries constitute the need for continued screening and development efforts.

### Resolution of inflammation: An active mechanism driven by newly discovered omega 3-fatty acid-derived mediators

The inflammatory response is divided into three temporal phases, initiation, amplification and maintenance, and resolution. Classical anti-inflammatory treatments have focused on interference with target pathways involved in the initiation and maintenance of inflammation. These strategies have only been partially successful, due to the large variety of pathways and pathologies involved. Moreover, selective and non-selective cyclooxygenase-2 (COX2) inhibitors show beneficial therapeutic effects but with obvious side effects. For many inflammatory conditions with symptoms resembling the outcomes of chemical threat injuries (such as asthma, ARDS, or dermatitis), clinical trials using cytokine inhibitors, steroids or NSAIDs have resulted in mixed outcomes, suggesting continued demand for the development of new clinical strategies counteracting inflammation.

A new area of inflammation research has focused on the process of inflammation resolution (29-32). Resolution of inflammation was thought to occur due to the lack of inflammatory drive when concentrations of inflammation-initiating and -maintaining mediators are removed or in decline. However, new studies show that the resolution of inflammation is an active mechanism involving the activation of signaling pathways during inflammation initiation, and the later generation of fatty-acid-derived specialized pro-resolving mediators (SPMs) that activate resolution mechanisms. These mediators include resolvins, lipoxins, protectins, and maresins. Resolvins and protectins are omega-3 fatty acids derived from docosahexaenoic acid (DHA; C22:6) or eicosapentaenoic acid (EPA; C20:5), the two dietary fatty acids widely consumed due to their reported anti-inflammatory and other beneficial effects (29). These mediators are produced enzymatically in human blood by neutrophils, with increased concentrations observed following aspirin treatment (33). SPMs are shown to bind to a range of G-protein-coupled receptors and ion channels such as transient receptor potential (TRP) ion channels to mediate their pro-resolving effects (29, 34, 35). In general, SPMs help injured tissues to return to their original architecture and function by promoting resolution through recruitment of non-inflammatory monocytes. Then, cellular debris, inflammation causative agent(s) and excess neutrophils will be removed by macrophages (30, 32, 36, 37).

### Therapeutic potential and clinical development of pro-resolving mediators

SPMs, when administered exogenously, have shown potent anti-inflammatory effects in a wide range of animal disease models, including lung injury, pain, dermatitis, arthritis, and colitis (32, 33, 35, 38-47). The development of SPMs as therapeutics is an active area of pharmaceutical research, with some SPMs such as Resolvin E1 and Resolvin D1 are currently in tests in clinical trials for the treatment of inflammatory eye conditions, psoriasis, chronic pain, and many others (https://clinicaltrials.gov/ct2/results?cond=&term=specialized+pro-resolving+mediators+OR+RvD1+OR+RvD2+OR+RvE1+OR+MaR1&cntry=&state=&city=&dist=&Se arch=Search, accessed 02/05/2019). In addition to bio-identical agents, chemically modified derivatives have been synthesized with improved stability and specificity (32, 36).

We hypothesize that accelerating resolution phase of inflammation may attenuate the exaggerated cutaneous inflammatory response following chemical threat agent exposure to sulfur mustard gas and tear gas agent, leading to decreased morbidity and improved recovery. Several new families of lipid-derived, local acting chemical mediators, which are derivatives of omega-3 fatty acids that enhance the resolution of the inflammatory process and restore the original architecture of injured tissues have been identified. The objective of these studies was to test the therapeutic potential of SPMs in mouse models of CS tear gas agent- and CEES-induced cutaneous injury.

## Materials and methods

Specialized pro-resolving mediators (Resolvin D1 (RvD1) and Resolvin D2 (RvD2)) were purchased from Cayman chemicals Inc., Ann Arbor, MI, USA. IRDye 800 CW was purchased from Li-Cor, Lincoln, NE, USA. 2-chloroethyl-ethyl-sulfide (CEES), a sulfur mustard analog was purchased from Sigma-Aldrich, MO, USA. CS tear gas agent was purchased from Combi-Blocks, San Diego, CA. All other chemicals and reagents used were obtained from scientific suppliers, such as Fisher Scientific and Sigma Aldrich.

C57BL/6 mice (male, 8 weeks) were purchased from Charles River, CT. Mice were housed in Association for Assessment and Accreditation of Laboratory Animal Care (AAALAC) International certified facilities. All animal protocols were approved by the Institutional Animal Care and Use Committee (IACUC), Yale University, New Haven, CT and Duke University School of Medicine, Durham, NC. Mice were given access to *ad libitum* mouse chow and water. Mice were given at least 48 hr acclimation period before initiating studies. In adherence to the Office of Animal Welfare Assurance guidelines, cage enrichment was done when mice were housed alone transiently.

### 2-chloroethyl-ethyl-sulfide mouse ear vesicant model (CEES MEVM)

Sulfur mustard analog-induced mouse ear vesicant model was developed as described in Achanta et al., 2018 (6). Briefly, the right ears of male 8 week old C57BL/6 mice were exposed to a total single dose of 0.2 mg (in 20 µL, ten microliters on each side of right ear) of 2-chloroethyl-ethyl-sulfide (CEES), a sulfur mustard analog following sevoflurane anesthesia. The left ears were exposed to vehicle (dichloromethane) that served as controls. One hour later, mice were administered a single dose of SPM (RvD1 or RvD2) at 2 µg/kg intraperitoneally (i.p), in PBS with 0.1% ethanol (injection volume: 10 mL/kg body weight). Control animals received vehicle (PBS with 0.1% ethanol) only. Figure 1A depicts the CEES MEVM study paradigm and dosage regimen. We prepared fresh SPM solutions within 30 minutes before injection. Briefly, SPMs were evaporated under a gentle stream of nitrogen gas and reconstituted in 0.1% ethanol in PBS.

**Figure 1.**
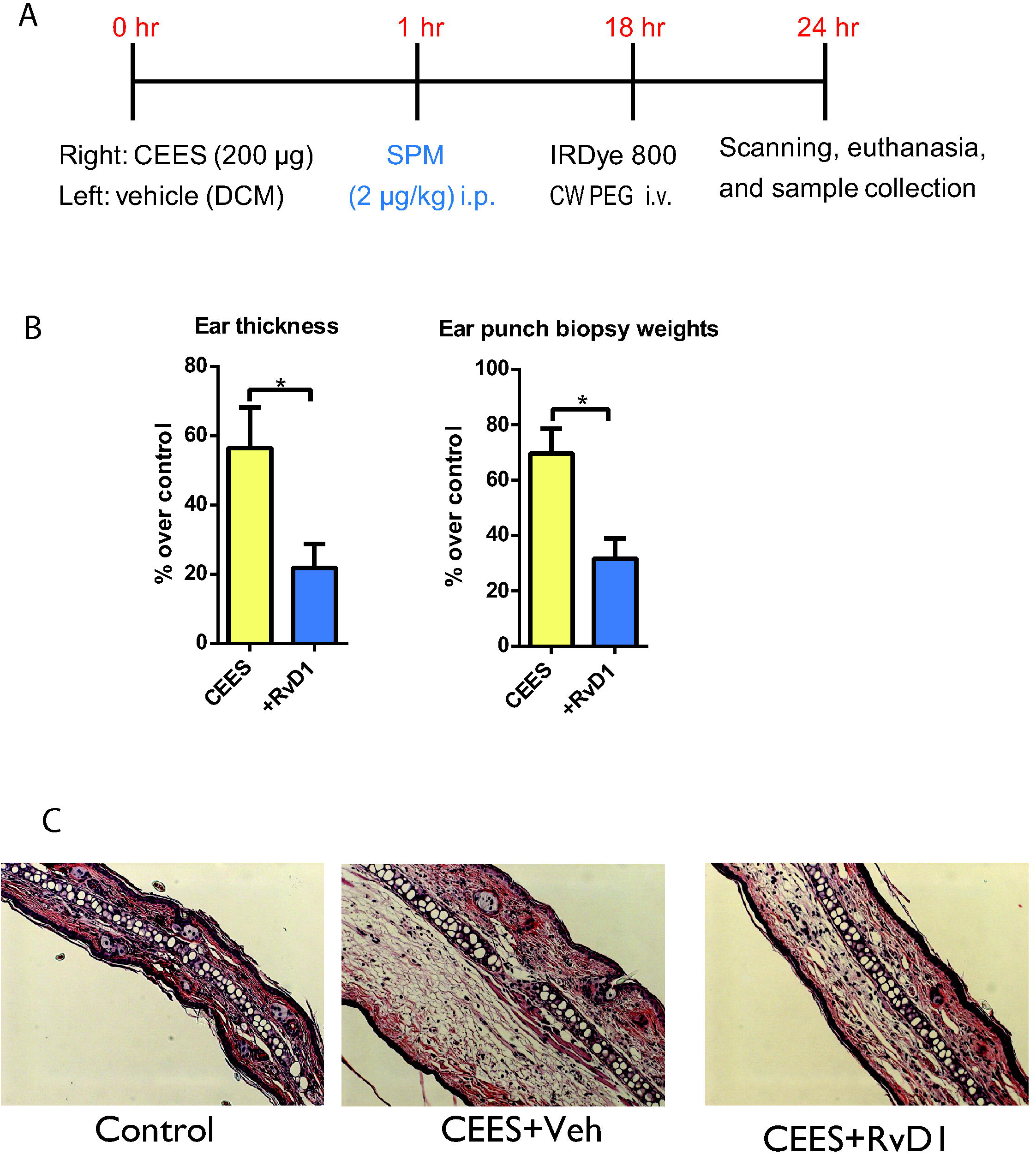
Gross morphological parameters and histopathology in CEES-exposed mice, treated with Resolvin D1 (RvD1) or vehicle. A) *2-chloroethyl-ethyl-sulfide mouse ear vesicant model (CEES MEVM) and treatment regimen*. Right and left ears of mice were exposed to CEES and vehicle (dichloromethane), respectively. RvD1 (2 µg/kg i.p) or vehicle was administered at 1 h post-CEES exposure. B) *Ear thickness and ear punch biopsy weights*. Ear thickness was measured by spring-loaded calipers. Ear punch biopsy weights were measured from three 4 mm circular ear punch biopsy samples. Percent decrease or increase in ear thickness and ear punch biopsy weights over control is presented. C) Representative H&E stained ear punch biopsy histopathology profiles are presented. Data are presented as mean ± SEM. n= 5/group.

### Visualizing extravasation of inflammatory exudate

To visualize the extravasation of inflammatory exudate into surrounding tissues and the healing process in CEES exposed ears, we injected IRDye 800CW intravenously at 18 hr post-CEES exposure under a brief sevoflurane anesthesia. We scanned anesthetized mice (ketamine and xylazine at 80-120 and 5-10 mg/kg body weight i.p., respectively) in dorsal recumbency at 23 hr post-CEES exposure using Li-Cor Odyssey CLX^®^ fitted with an accessory, Mouse-Pod^®^ (Li-Cor, Nebraska, USA). Mice were maintained at 37 °C while under anesthesia and during the scanning procedure. We used the following settings for scanning of mice - channels: 700/800; scan resolution: 42 µm, intensities: 1 for both channels; data analysis: small animal; scan quality: medium (increasing the quality of scan will increase the time of scanning); focal offset: 1 mm. Data were analyzed using the Small Animal Image Analysis Suite provided by the manufacturer.

### Ear punch biopsy sample collection, pro-inflammatory cytokine, and histopathology analysis

Mice were euthanized 24 hr post-CEES exposure in the CO_2_ chamber and then by a secondary method, approved by the IACUC and AVMA guidelines for euthanasia. Ear thickness was measured using spring-loaded electronic calipers (Mitutoyo QUICKmini, Japan). Three 4 mm ear punch biopsies were excised (4 mm Biopsy Punch, Miltex Inc., York, PA, USA) and weighed to determine edema as described previously (6). One punch biopsy was used to determine concentrations of pro-inflammatory mediators known to contribute to the vesicant injury. Briefly, ear punch biopsy samples were homogenized in lysis buffer (50 mM Tris-base, 150 mM NaCl, 5 mM EGTA supplemented with EDTA-free complete protease inhibitor (Roche Diagnostics GmbH, Mannheim, Germany) and 0.5% Triton X-100), using a Bullet Blender and Zirconium Oxide Beads (NextAdvance^®^, Averill Park, NY) (6). Using the enzyme-linked immunosorbent assay (ELISA), we examined the concentrations of matrix metalloproteinase 9 (MMP-9), IL-1β (interleukin-1 beta), KC/CXCL1 (keratinocyte chemoattractant)/(chemokine(C-K-X motif) ligand 1), MIP-2/CXCL2 (macrophage inflammatory protein 2)/(chemokine(C-K-X motif) ligand 2), and interleukin-6 (IL-6) in homogenized supernatant protein extracts of ear punch biopsy samples. R&D Systems cytokine kits (Minneapolis, MN) or a high throughput multiplex cytokine assay system (Milliplex MAP Mouse Cytokine/Chemokine Magnetic Bead Panel, Millipore, MO, USA) was used for cytokine quantification, following manufacturer’s instructions. All samples were analyzed in at least duplicate on Infinite M200 Pro (Tecan, Germany) or Bio-Plex 200 system (Bio-Rad, Hercules, CA). The concentrations of cytokines were quantified using a standard curve or a 5-parameter logistic regression analysis. Concentrations of samples outside the range of standard curves were excluded. The concentrations of protein in homogenate samples were determined using Pierce BCA protein assay (Thermo Scientific, Rockford, IL).

One punch biopsy was fixed in 10% formaldehyde, embedded in paraffin, sectioned at 5 µm thickness, and stained with hematoxylin and eosin (H&E) as per standard protocols. Images were obtained with a Zeiss Axio Imager Z1 microscope and analyzed by AxioVision Rel. 4.7 software (Zeiss, Munich, Germany). Cutaneous histopathology was assessed based on guidelines in Silny et al., 2005 (6, 48).

### Electrophilic agent (CS tear gas agent) induced cutaneous inflammation

Mice were briefly anesthetized with sevoflurane. We exposed right ears of mice with 20 µL (10 µL on each surface) of 200 mM CS tear gas agent and applied an equal volume of dimethyl sulfoxide (DMSO, a vehicle for CS tear gas agent application) on left ears. We administered either SPM (RvD1 or RvD2) at 5 µg/kg body weight i.p or vehicle (PBS with 0.1% ethanol) at 30 minutes and 4 hours post-CS exposure. We euthanized mice at 6.5 hours post-CS exposure. Ear thicknesses, ear punch biopsy weights; tissue homogenization to harvest protein extracts, cytokine analysis, and H&E staining were done as described in the above CEES MEVM sections. Figure 5A depicts the study paradigm and dosing regimen.

**Figure 2.**
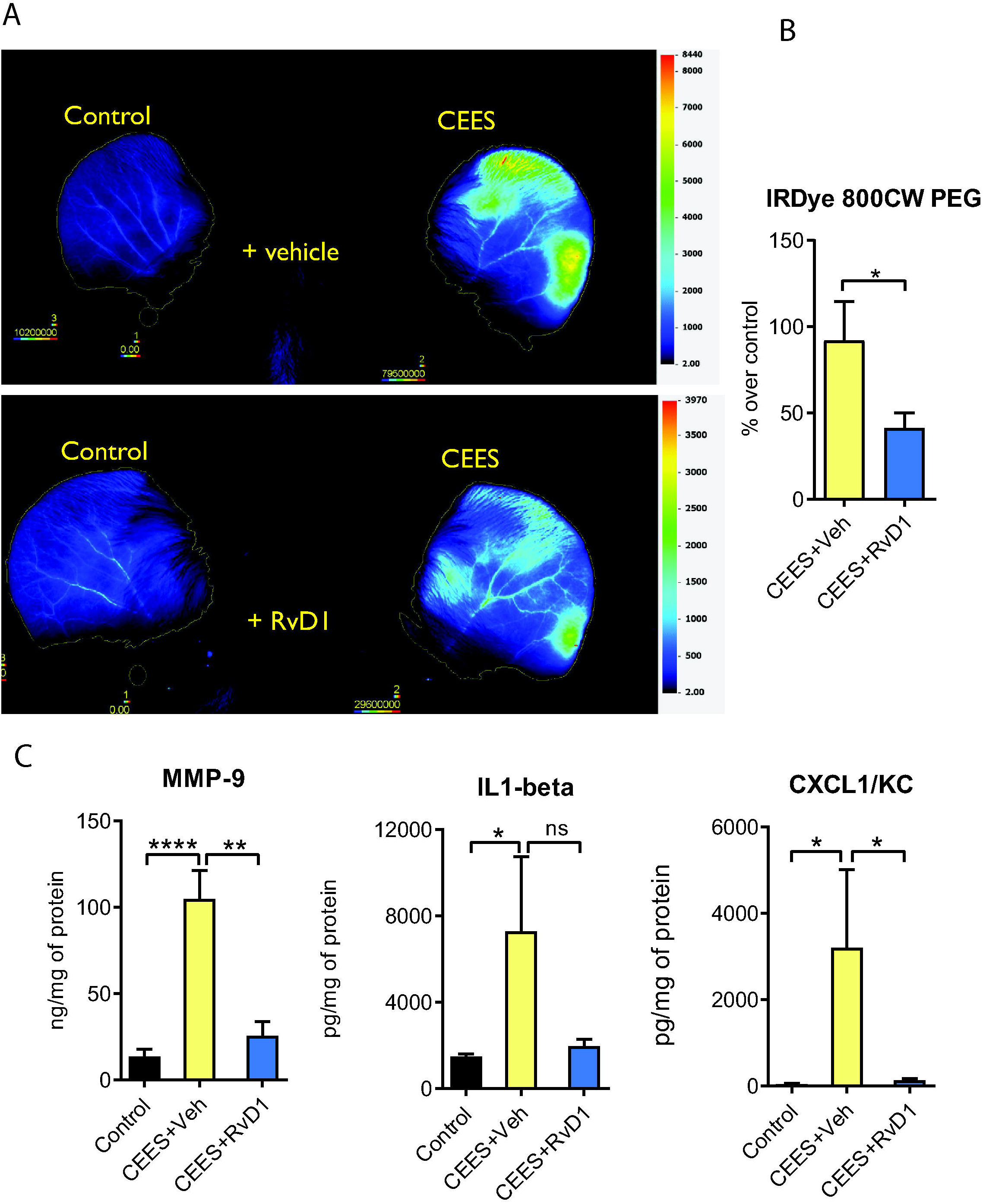
Therapeutic effects of Resolvin D1 on vascular leakage and pro-inflammatory cytokines in ear punch biopsy homogenates of CEES-exposed mice. 2-chloroethyl-ethyl-sulfide mouse ear vesicant model (CEES MEVM) and treatment regimen are as in Figure 1A. A) *Representative images of vascular leakage in CEES MEVM*. Extravasation of inflammatory exudate into surrounding tissues and the resolution of inflammation with the administration of RvD1 or vehicle in CEES-exposed mice was studied by injecting IRDye 800CW intravenously at 18 h post-CEES exposure. Mice were scanned at 23 h post-CEES exposure using Li-Cor Odyssey CLX^®^. B) The percent increase or decrease in intensity of dye in the CEES-exposed ear over control ear is presented (n=5/group). C) Expression of MMP-9 (n=10-11/group) and pro-inflammatory cytokines such as IL-1β (5-9/group) and CXCL1/KC (4-8/group) in ear punch biopsy homogenate samples using ELISA. Data are presented as mean ± SEM.

**Figure 3.**
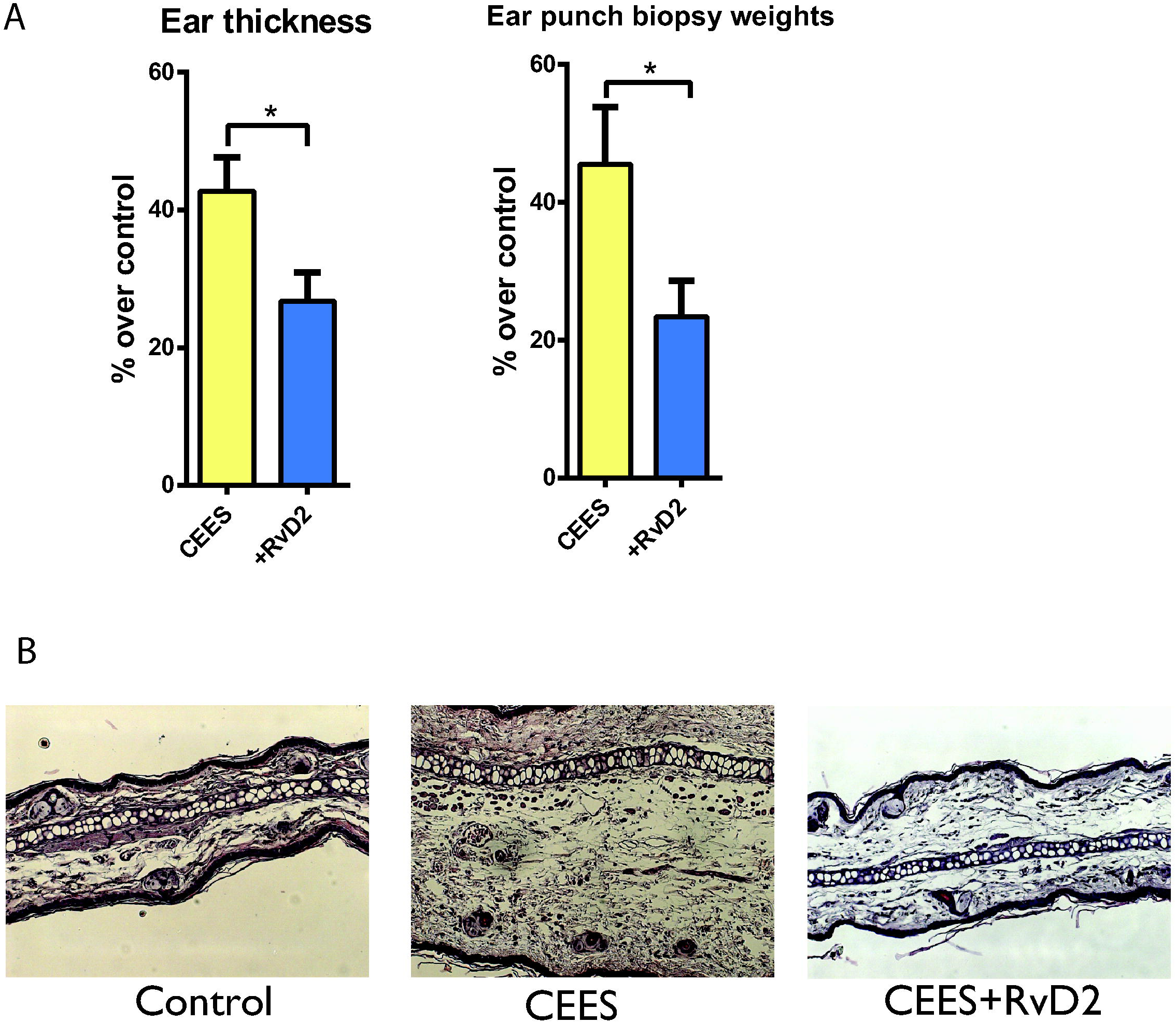
Gross morphological parameters and histopathology in CEES-exposed mice, treated with Resolvin D2 (RvD2) or vehicle. 2-chloroethyl-ethyl-sulfide mouse ear vesicant model (CEES MEVM) and treatment regimen are as in Figure 1A. Mice received either Resolvin D2 (RvD2) at a dose rate of 2 µg/kg i.p body weight or vehicle 1 h post-CEES exposure. A) Ear thickness and ear punch biopsy weights. Ear thickness was measured using spring-loaded calipers. Ear punch biopsy weights were determined from three 4 mm ear punch biopsy samples from each ear. Percent increase or decrease in ear thickness and ear punch biopsy weights over control are presented. B) Representative H&E stained ear punch biopsy histopathology samples are presented. Data are presented as mean ± SEM. n = 10/group.

**Figure 4.**
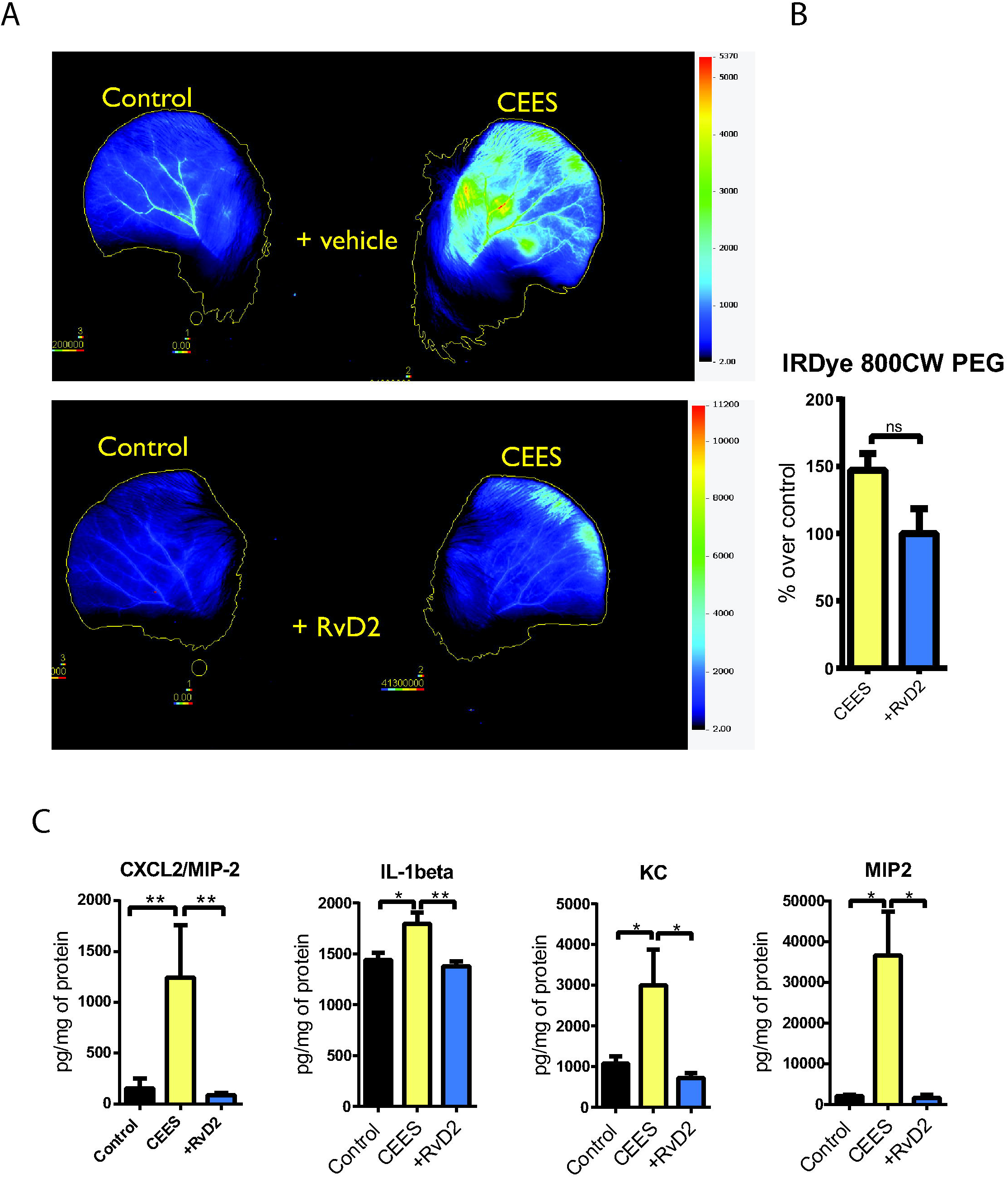
Therapeutic effects of Resolvin D2 on vascular leakage and pro-inflammatory cytokines in ear punch biopsy homogenates of CEES-exposed mice. 2-chloroethyl-ethyl-sulfide mouse ear vesicant model (CEES MEVM) and treatment regimen are as in Figure 1A. A) *Representative images of vascular leakage in CEES MEVM*. Extravasation of inflammatory exudate into surrounding tissues and the resolution of inflammation with the administration of RvD2 or vehicle in CEES-exposed mice was studied by injecting IRDye 800CW intravenously at 18 h post-CEES exposure. Mice were scanned at 23 h post-CEES exposure using Li-Cor Odyssey CLX^®^. B) The percent increase or decrease in the intensity of dye in the CEES-exposed over control ears is presented (n=3-4/group). C) Expression of pro-inflammatory cytokines such as CXCL2/MIP-2, IL-1β and CXCL1/KC in ear punch biopsy homogenate samples using ELISA (n=3-6/group). Data as mean ± SEM.

**Figure 5.**
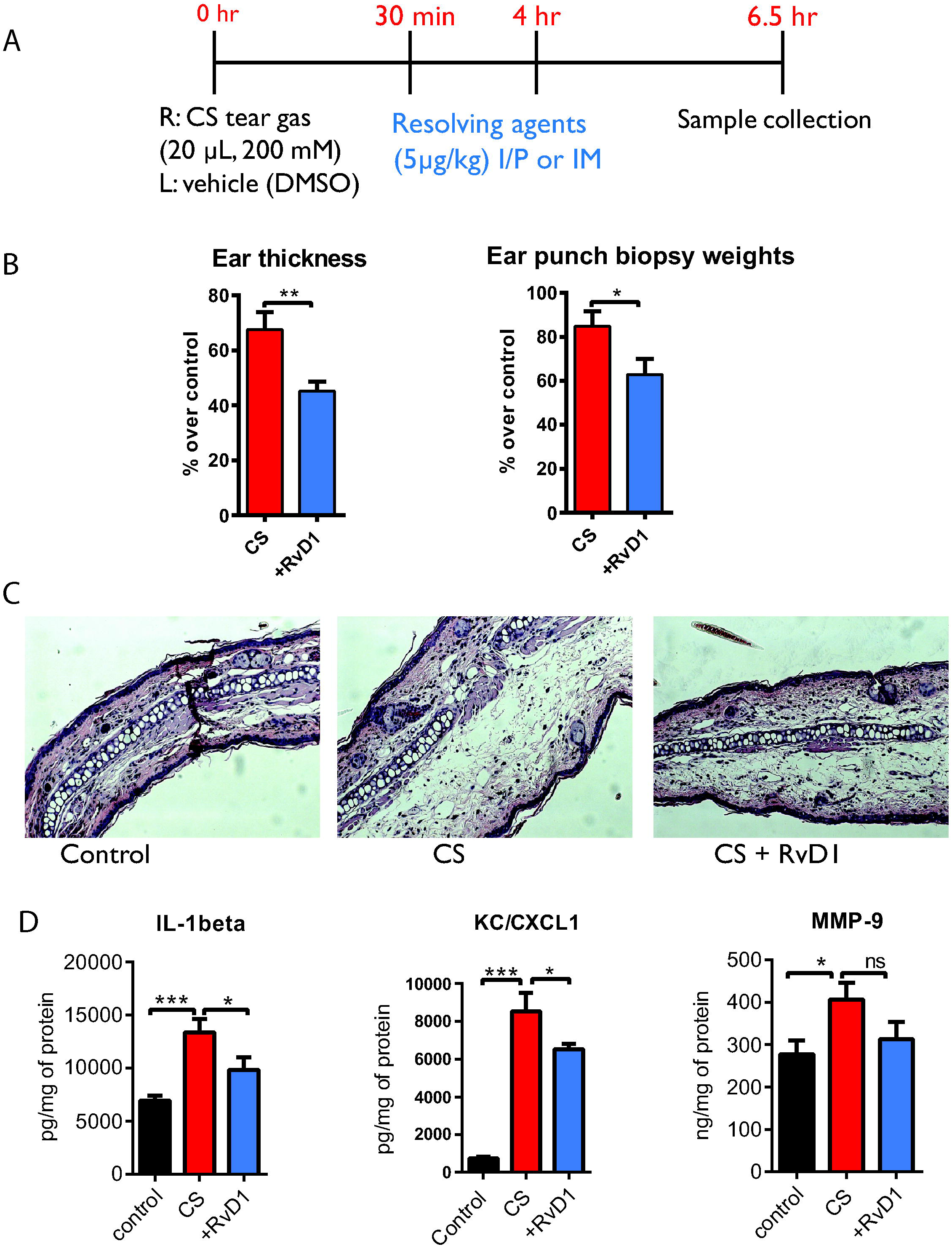
Therapeutic effects of Resolvin D1 (RvD1) in CS tear gas-induced acute skin inflammation. A) CS tear gas exposure and treatment regimen. Right and left ears of mice were exposed to CS tear gas agent and vehicle (DMSO), respectively. Mice received either RvD1 at a dose rate of 5 µg/kg body weight at 30 minutes and 4-hour post-CS tear gas exposure or vehicle (0.1% ethanol in PBS) intraperitoneally. B) Ear thickness and ear punch biopsy weights. Ear thickness was measured by spring-loaded calipers. Ear punch biopsy weights were measured from three 4 mm circular ear punch biopsy samples. Percent decrease or increase in ear thickness and ear punch biopsy weights over control is presented (n=9-10/group). C) Representative H&E stained ear punch biopsy histopathology samples are presented. D) Expression of MMP-9 and pro-inflammatory cytokines such as IL-1β and CXCL1/KC in ear punch biopsy homogenate samples using ELISA (n=3-8/group). Data are presented as mean ± SEM.

### Data Analysis and Statistics

Data were analyzed using GraphPad Prism 7 for Windows, GraphPad Software, La Jolla, CA, USA. Statistical difference was tested either by an unpaired two-tailed t-test or by one-way ANOVA with Tukey’s Multiple Comparison Test. Error bars are represented as the standard error of mean estimate (SEM). Statistical significance was denoted by *p<0.05 or **p<0.01 or ***p<0.001.

## Results

### CEES MEVM

Cutaneous exposure to CEES resulted in tissue edema and microvesicles. Ear thickness, measured with spring-loaded calipers, increased by 49.4±21% compared to vehicle (dichloromethane) exposed ears. Ear punch biopsy weights, measured from three 4-mm punch biopsies, increased by 57.5±23% compared to vehicle exposed ears. Treatment of mice with RvD1 or RvD2 post-CEES exposure decreased tissue edema, measured with ear thickness and ear punch biopsy weights (Figures 1B and 3A). Profound vascular leakage was noted in ears exposed to CEES due to vascular dilation, increased diapedesis, and accumulation of inflammatory exudate in the subcutaneous tissue. Whereas, vascular leakage was significantly decreased in mice that were treated with RvD1 or RvD2 post-CEES exposure (Figures 2A-B and 4A-B). The intensity of IRDye 800 CW in SPM-treated groups was significantly lower than the vehicle-treated animals. Inflammatory cytokine markers, such as MMP-9, IL-1β, CXCL1/KC, CXCL2/MIP2, and IL-6, were decreased in treatment groups compared to control (vehicle) mice, suggesting that acceleration of inflammation resolution resulted in diminished tissue destruction by proteases and recruitment of inflammatory neutrophils (Figures 2C and 4C). Tissue biopsies of CEES exposed mice treated with SPMs showed dramatically attenuated edema and microvesicle formation (separation of epidermal and dermal layers) and reduced inflammatory infiltrating cells, resulting in almost normal ear thickness (Figures 1C and 3B).

### CS tear gas

Exposure to CS tear gas agent-induced acute inflammation, evidenced by erythema and edema within few minutes after exposure. Compared to the vehicle (DMSO-exposed ears), ear thickness and ear punch biopsy weights increased by 63±21% and 75±23%, respectively.

Post-exposure treatment of CS tear gas agent-induced acute cutaneous inflammation with SPMs (RvD1 or RvD2) decreased tissue edema compared to the control group (Figure 5B and 6A). SPMs decreased pro-inflammatory cytokines (IL-1β, CXCL1/KC, CXCL2/MIP2, and IL-6) and MMP-9 (Figures 3B and 4B). In tissue histopathology sections of CS tear exposed ears, edema was an obvious finding in addition to infiltration of cells. In treated groups, edema, infiltration of pro-inflammatory cells, and keratinolysis were decreased compared to control groups (Figure 3C and 4C).

**Figure 6.**
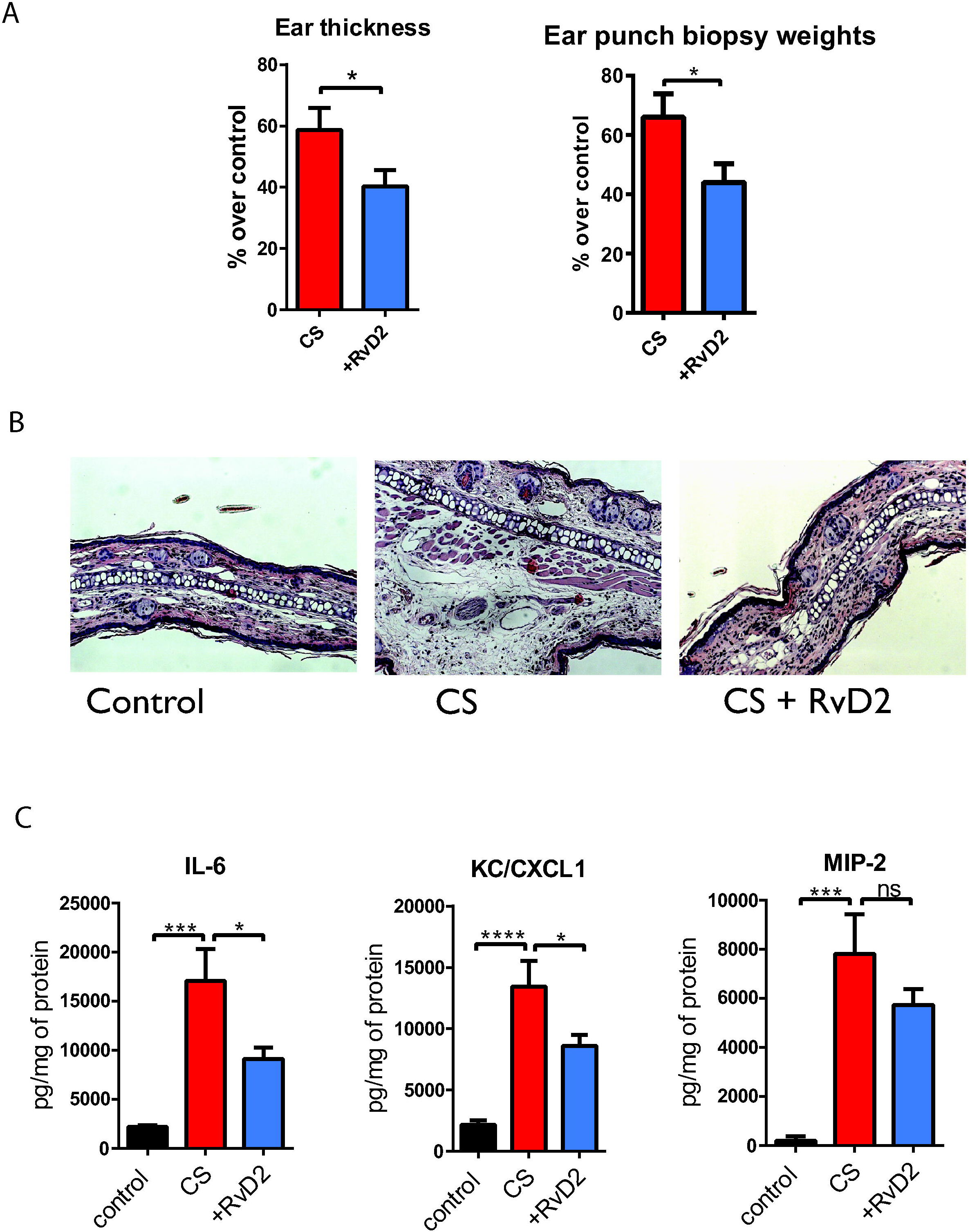
Therapeutic effects of Resolvin D2 (RvD2) in CS tear gas-induced acute skin inflammation. CS tear gas exposure and treatment regimen are as in Figure 5A. Mice received either RvD2 at a dose rate of 5 µg/kg body weight at 30 minutes and 4-hour post-CS tear gas exposure or vehicle (0.1% ethanol in PBS) intraperitoneally. A) Ear thickness and ear punch biopsy weights. Ear thickness was measured by spring-loaded calipers. Ear punch biopsy weights were measured from three 4 mm circular ear punch biopsy samples. Percent decrease or increase in ear thickness and ear punch biopsy weights over control is presented (n=10/group). B) Representative H&E stained ear punch biopsy histopathology samples are presented. C) Expression of pro-inflammatory cytokines such as IL-6, CXCL1/KC, and MIP-2 in ear punch biopsy homogenate samples using ELISA (n=4-6/group). Data are presented as mean ± SEM.

## Discussion

Skin is the body’s largest organ system with an average body surface area ranging from 1.6 – 1.9 m^2^ in humans. Therefore, cutaneous injuries following chemical exposure are the most common sequelae. In this study, skin injury caused by sulfur mustard analog, CEES and CS tear agent were studied in mice; and the therapeutic potential of SPMs was investigated in chemical threat agent-induced cutaneous injuries. To the best of authors’ knowledge, this is the first report on the use of SPMs as therapeutic agents in chemical agent-induced cutaneous inflammation or in any other medical countermeasure studies.

### CEES MEVM

Sulfur mustard interacts with biological matrices in a wide variety of cascade of pathophysiological events and therefore, difficult to control inflammation. Although the toxicological effects of sulfur mustard on the ocular and respiratory system are widely studied, information for cutaneous effects is scarce. CEES is commonly used as an analog to study the effects of sulfur mustard (HD). CEES MEVM in this study recapitulated previously published reports elsewhere (5, 6). Edema or erythema of exposed ears was not seen immediately. CEES is a mono-functional alkylating analog of sulfur mustard which forms adducts with the biological interface, unlike sulfur mustard. CEES is 100 times less potent than sulfur mustard. Visible blisters/vesicles were not seen in CEES MEVM, unlike sulfur mustard. However, CEES is commonly used as a surrogate agent to study the effects of sulfur mustard under laboratory settings due to its similarities of chemical and toxicological effects to sulfur mustard.

In this study, several pro-inflammatory cytokines markers were tested, such as MMP-9, IL-1β, CXCL1/KC, CXCL2/MIP-2, and IL-6. MMP-9 is thought to be a major effector of tissue destruction caused by vesicants and may contribute to blister formation by separating the dermis from the epidermis (49, 50). KC is a potent neutrophil attractant, guiding pro-inflammatory neutrophils into the injured tissue. IL-1β is produced by macrophages and various other cell types in response to inflammatory agents, infections, or microbial endotoxins. IL-1β is not produced in healthy tissues, with the exception of skin keratinocytes, some epithelial cells, and certain cells of the nervous system. In our studies, we observed similar findings that support the expression of IL-1β in control ears. In these studies, although inflammatory cytokine makers were decreased in treatment groups compared to the vehicle group, some variability was observed between SPMs tested.

We developed an *in vivo* imaging with infra-red fluorescent dye to visualize extravasation of inflammatory exudate and real-time healing. Although such imaging is done frequently in tumor biology studies, this is the first of its kind *in vivo* imaging technique in cutaneous inflammatory studies (51). Increased vascular leakage was noted in mouse ears exposed to CEES whereas treatment with SPMs decreased vascular leakage. Vascular leakage was quantified with the intensity of fluorescence. Treatment with SPMs significantly decreased fluorescence of IRDye 800 CW PEG. Vascular leakage is a common feature of inflammation. Vascular permeability is a result of increased capillary hydrostatic pressure and increased spaces between vascular endothelial linings. The non-specific fluorescent dye, IRDye 800CW PEG contrast agent (25-60 kDa), used in this study reaches the sites of acute inflammation due to vascular leakage and decreased return to the vasculature due to alteration in colloid osmotic pressure. We visualized ear vasculature in mice as early as 10 minutes post-administration of dye. The method was optimized to track the progression of inflammation in real time with this less invasive procedure. In this study, we injected IRDye 800CW PEG agent at 1:2 dilution (0.5 nmol per animal) instead of the manufacturer’s suggested amount (1 nmol per animal) to avoid signal saturation. IRDye 800 CW is excreted in urine. Therefore, temporary individual caging of mice with cage enrichment after the injection of dye is suggested to prevent background noise from urine excreted by the cage mates. This technique is highly useful in tracking the therapeutic efficacy of potential drug candidates in long-term skin injury studies.

### CS tear gas

In recent years, the deployment of CS tear gas agent has significantly increased worldwide, including the United States for riot control. Although tear gas agents have been used since several decades for riot control and to incapacitate enemies in war fronts, no specific therapeutic agents are available for treating injuries caused by tear gas agents. In a real riot control situation, the actual concentration of exposure of CS tear agent is unknown and is highly variable. In our *in vivo* CS tear gas agent exposure titration studies, we found that exposure to concentrations of 50 - 200 mM of CS tear gas agent saturated the response. However, in this study, we tested the therapeutic potential of pro-resolving agents at 200 mM concentration (20 µL) of CS tear agent to show therapeutic benefits even at higher CS tear gas exposure levels.

The clinical manifestation of CS tear gas exposure includes immediate burning and pain sensation. These signs are the result of intense sensory irritation at various body sites. Studies in our laboratory have shown that CS tear gas exerts its effect through the activation of TRPA1 channels (8, 52). After the removal of patients from CS tear gas exposure site, lacrimation and burning sensation may decrease within 30 minutes. However, exposure to higher concentrations of this xenobiotic or for a prolonged period will result in severe cutaneous injuries.

In this study, edema and erythema were the most significant findings of cutaneous injury upon exposure to CS tear gas agent. Both ear punch biopsy gross markers and histopathology demonstrated heightened edema with CS tear gas exposure. Treatment with SPMs significantly reduced inflammation. Similar anti-edematogenic effects of SPMs were observed in carrageenan-induced paw edema (53). Pro-inflammatory markers were significantly increased in CS tear gas agent exposed mice whereas treatment with pro-resolving agents decreased these cytokine markers.

### Dose and route of administration

The results from this study were promising in showing the therapeutic effects of SPMs in two chemical threat agents-induced cutaneous inflammation. There are several teams studying the potential use of these agents as therapeutics against several indications around the world (30, 32, 33, 36, 38, 39, 47, 54-57). Based on previous studies in other disease modalities in mouse models, nano-gram amounts were administered in these studies and these doses were shown to be therapeutically effective. In this study, therapeutic efficacies of pro-resolving agents were demonstrated in the intraperitoneal route of administration. However, in mass casualty situations, auto-injectors for intramuscular injection or oral medication would be ideal. Testing the efficacy of pro-resolving agents through other routes of administration, such as intramuscular and subcutaneous are warranted. It is expected that the intramuscular and subcutaneous route of administration would be more efficacious compared to the variable intraperitoneal route of administration.

In the continuum of these studies, testing therapeutic potential in other routes of administration, testing combination of pro-resolving agents to get the synergistic beneficial effects and testing in higher mammalian species are warranted. Additional pro-resolving agents are discovered recently and it’s worth to screen therapeutic potential of these new agents in chemical cutaneous injury models.

### Mechanism of action of SPM

Although the mechanism of action of pro-resolving agents in this study was not elucidated, the published literature shows several possible mechanisms of action (30, 55, 58). Recent studies show that pro-resolving agents work through G protein-coupled receptors (GPCRs), such as ALX/FPR2, GPR18 and GPR32, whereas a few other studies show that these agents work through inhibition of transient receptor potential channels (35, 42, 59-61). Achanta et al., Bessac et al., and Stenger et al., have shown the involvement of transient receptor potential ankyrin repeat 1 (TRPA1) ion channel in mediating the effects of CEES and CS tear gas agents using *in vitro* and *in vivo* studies (6, 52, 62-64). Park et al and others showed that RvD1 and RvD2 regulate TRPA1 currents in mouse DRG neurons (35, 59, 65). Based on the literature review, we believe that SPMs mitigated the cutaneous injury caused by CEES and CS tear gas agents through inhibition of TRPA1 ion channels (Figure 7). The interactions between GPCRs and TRP ion channels further substantiate our proposed mechanism of action of SPMs (RvD1 or RvD2) through inhibition of TRPA1 ion channels (66, 67). However, further *in vitro* studies with TRPA1 transfection in HEK293T cells or *in vivo* studies with TRPA1 knock mouse are warranted to test our interpretation.

**Figure 7.**
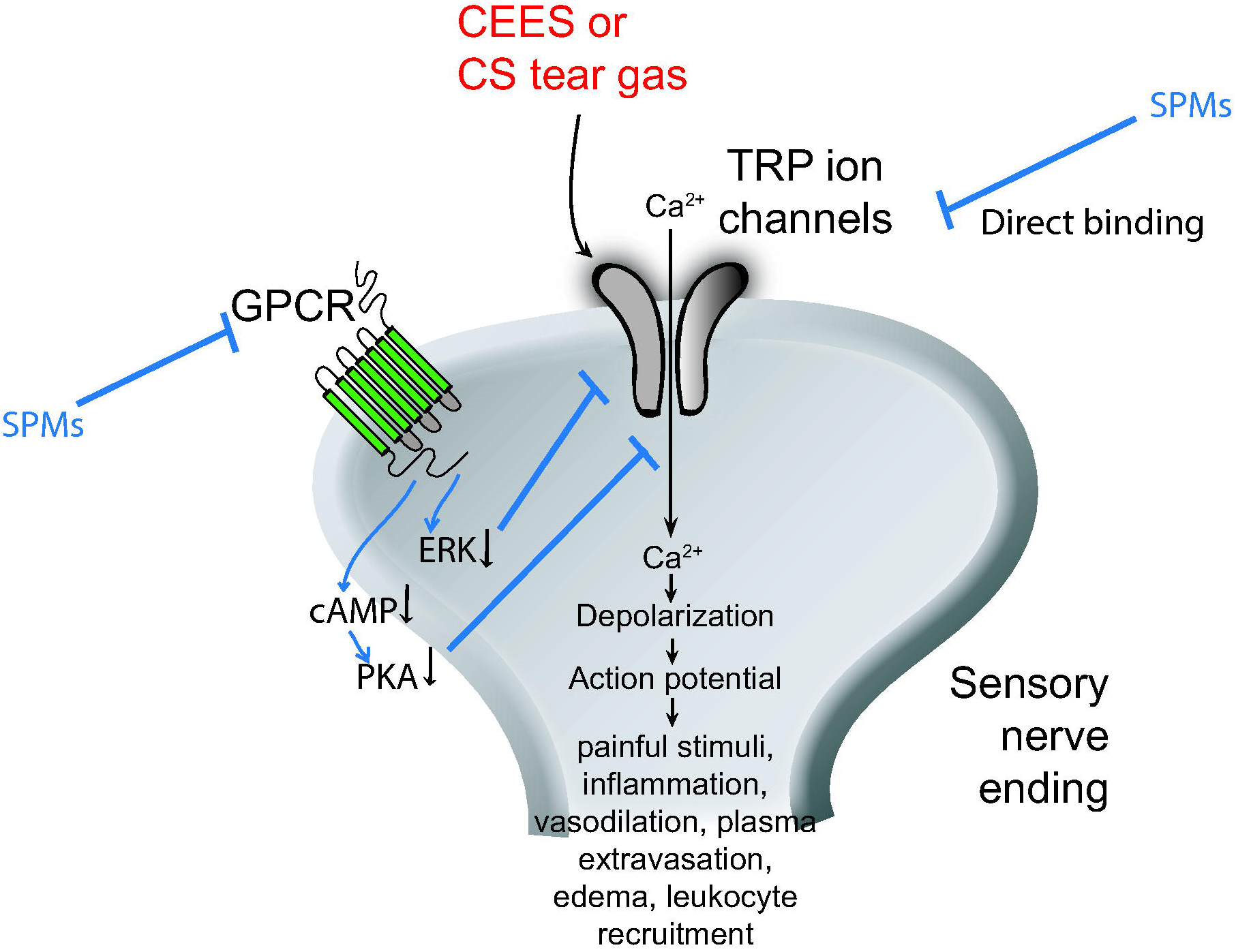
Mechanism of action of SPMs through GPCR-TRP ion channel axis. Transient receptor potential Ankyrin repeat 1 (TRPA1) ion channels mediate the effects of CEES and CS tear gas agents. SPMs (RvD1 or RvD2) ameliorate the inflammation caused by CEES and CS tear agents through inhibition of TRPA1 ion channels directly or through modulation of G-protein coupled receptors (GPCRs) which subsequently inhibits TRPA1 ion channels.

### Potential of pro-resolving agents as therapeutic candidates

Due to the similarity of SPMs to endogenous lipid mediators, and because of the low dosages required to initiate resolution of inflammation, it is expected that these agents will display minimal side effects or toxicity compared to the conventional NSAIDs or opioids. No side effects or toxicity of SPMs was published in the literature. A combination of these SPMs might also offer better therapeutic outcomes compared to single-agent therapy which is worth testing. Few of the limitations of SPMs are lack of stability at room temperature and non-availability of commercial SPMs as drug formulations. Some of the commercial nutraceuticals enriched with metabolites of EPA and DHA such as 17-hydroxy-docosahexaenoic acid (17-HDHA), 18-hydroxy-eicosapentaenoic acid (18-HEPE), and 14-DDHA might be worthwhile to test in the skin injury models for their efficacy (https://www.metagenics.com/spm-active, accessed 02/11/2019; http://www.solutex.es/en/products/biolipidos/lipinova, accessed 02/11/2019). If the therapeutic potential of these SPMs is established in rodent and non-rodent higher mammalian species, then the approval will be feasible for the indications of sulfur mustard and tear gas agent-induced cutaneous inflammation under the US FDA’s animal rule, unlike other new chemical entities (https://www.fda.gov/downloads/drugs/guidances/ucm399217.pdf, accessed 10/23/2018).

## Conclusion

Exposure of CS tear gas agent and CEES to ear skin caused profound cutaneous inflammation in mouse models. Post-exposure treatment with specialized pro-resolving mediators decreased inflammation. These SPMs (RvD1 and RvD2) can be used as potential medical countermeasures to hasten the resolution of inflammation caused by chemical threat agents.

## Acknowledgments

We appreciate thoughtful discussions of Drs. Bruce Levy, MD and Charles Serhan, PhD, Harvard University, Boston, MA, USA. This study was supported by grants from NIH (NIEHS): R21ES022875-01 (awarded to SEJ). The content is solely the responsibility of the author and does not necessarily represent the views of the NIH.

## Conflict of interest

Authors do not have any conflicts of interests to disclose

## Author contributions

SA, NC, and SEJ designed research; SA, NC, BL, and SB performed research; SA and NC conducted *in vivo* studies independently. SA developed and optimized new *in vivo* imaging with an infrared fluorescent dye; SA, NC, and SEJ analyzed data; SA wrote the first draft of manuscript; SA and SEJ approved the final version.

## Abbreviations

RvD1: (Resolvin D1);
RvD2: (Resolvin D2);
CS: (2-Chlorobenzalmalononitrile);
CEES: (2-chloroethyl-ethyl-sulfide);
SM: (sulfur mustard);
HD: (sulfur mustard);
MMP-9: (matrix metallo protein-9),
IL-1β: (Interleukin-1 beta),
KC/CXCL1: (keratinocyte chemoattractant)/(chemokine (C-X-C motif) ligand 1);
CXCL2: (chemokine (C-X-C motif) ligand 2);
IL-6: (interleukin-6);
ELISA: (enzyme linked immune sorbent assay);
PBS: (phosphate buffer saline);
MEVM: (mouse ear vesicant model);
i.p: (intraperitoneal);
i.v: (intravenous);
NSAIDs: (nonsteroidal anti-inflammatory drugs)

